# VariPred: Enhancing Pathogenicity Prediction of Missense Variants Using Protein Language Models

**DOI:** 10.1101/2023.03.16.532942

**Authors:** Weining Lin, Jude Wells, Zeyuan Wang, Christine Orengo, Andrew C.R. Martin

**Affiliations:** Institute of Structural and Molecular Biology, Division of Biosciences, University College London, London, United Kingdom; Department of Computer Science, University College London, London, United Kingdom; College of Computer Science and Technology, Zhejiang University, Zhejiang, China

**Author notes:** Corresponding author. Christine Orengo & Andrew C.R.Martin, Institute of Structural and Molecular Biology, Division of Biosciences, University College London, United Kingdom. or.

**Keywords:** VariPred, Bioinformatics Tool, Clinical Significance Database, Protein Language Model, Data Analysis, Variant Impact Prediction

## Abstract

Computational approaches for predicting the pathogenicity of genetic variants have advanced in recent years. These methods enable researchers to determine the possible clinical impact of rare and novel variants. Historically these prediction methods used hand-crafted features based on structural, evolutionary, or physiochemical properties of the variant. In this study we propose a novel framework that leverages the power of pre-trained protein language models to predict variant pathogenicity. We show that our approach VariPred (**Var**iant **i**mpact **Pred**ictor) outperforms current state-of-the-art methods by using an end-to-end model that only requires the protein sequence as input. By exploiting one of the best performing protein language models (ESM-1b), we established a robust classifier, VariPred, requiring no pre-calculation of structural features or multiple sequence alignments. We compared the performance of VariPred with other representative models including 3Cnet, EVE and ‘ESM variant’. VariPred outperformed all these methods on the ClinVar dataset achieving an MCC of 0.751 *vs*. an MCC of 0.690 for the next closest predictor.

## 1 Introduction

A large portion of genetic variation is represented by single nucleotide variants (SNVs). SNVs occur in both protein coding and non-coding regions, while protein-coding SNVs can be further divided into synonymous and non-synonymous (nsSNVs) types. Synonymous SNVs do not change the amino acid sequence of the resulting protein while non-synonymous SNVs (nsSNVs) do.

Missense mutations, in which a single amino acid is replaced by another, are the most common type of nsSNV (the others leading to truncation or extension). There is a long history of using physicochemical and evolutionary information to predict whether a given missense mutation is disease-causing [1–3]. Nonetheless it remains a major challenge to predict pathogenicity.

To tackle these challenges, many computational tools based on supervised machine learning techniques have been developed to predict the potential impact of variants. These compute deleterious scores based on dozens of biological properties of variants, such as evolutionary conservation [3–5], biochemical properties of amino acids [6,7], and structural features of proteins [8,9].

However, typically only a subset of variants can be annotated with all the features. This is especially true for tools that require a protein structure. There are 200 million protein sequences in the UniProt databank dated 12^th^ Oct 2022 (see: https://www.ebi.ac.uk/uniprot/TrEMBLstats), but only 200,000 experimentally-determined protein 3D structures stored in the Protein Data Bank (see: https://research.rutgers.edu/news/new-collaboration-between-rcsb-protein-data-bank-and-amazon-web-services-provides-expanded). This indicates that only approximately one in a thousand proteins have a reliable, experimentally resolved structure. For example, a commonly used predictor, Missense3D, can only structurally annotate 1965 and 2134 variants out of 26,884 disease-associated and 563,099 neutral variants, using structures from the PDB [9]. Even given the increase in structural coverage using predicted protein structures from AlphaFold2 [10], the accuracy of AlphaFold2 in predicting the structure of proteins with shallow multiple sequence alignments (MSAs) or orphan proteins, is questionable [11] and structure-based predictors may need to be re-trained for different levels of predicted quality obtained from AlphaFold2.

One downside of supervised machine learning models is that they can be prone to overfitting. This problem is particularly pronounced in cases where the training data contains genes where all variants have the same class label (benign / pathogenic). A previous study identifies this as the ‘Type 2 data circularity’ problem [12]. To avoid these problems, unsupervised learning models such as EVE [13] were developed. EVE makes predictions using features derived from MSAs. At the time of EVE’s release the authors reported state-of-the-art performance (AUC-ROC score of 0.91) for proteins that have been associated with disease in the clinical database ClinVar [13]. In a comparison between EVE and other computational variant effect predictors, EVE outperforms most widely used tools, including PolyPhen-2 [6], SIFT [14] and CADD [15] which are the three most popular predictors. However, EVE’s prediction accuracy for proteins not covered by informative MSAs remains unexplored.

In the latest Critical Assessment of Genome Interpretation (CAGI-6), a novel predictor named 3Cnet was top ranked in the SickKids clinical genomes and transcriptomes panel (see: https://www.3billion.io/blog/3billion-wins-in-cagi6-a-global-artificial-intelligence-genome-interpretation-contest/). 3Cnet is a deep artificial LSTM-based neural network model, which utilizes multiple protein features including MSAs, amino acid physicochemical properties, and features such as motifs and active sites as the input [16]. As EVE and 3Cnet are both trained for predicting the clinical significance of missense variants, and both have reported state-of-the-art performance in this field, we have selected these as state-of-the-art methods against which to benchmark our approach.

Most recent novel protein data-representation approaches take inspiration from language models that have yielded ground-breaking improvements in natural language processing (NLP). In particular, the Transformer neural network architecture [17], can learn contextualised word representations from a large amount of unlabelled text data and has achieved state-of-the-art performance for several NLP tasks. In the life sciences, most protein language models (PLMs) use Transformer architectures which were developed for NLP, but were subsequently trained on protein sequences with the goal of deciphering the ‘natural language’ of proteins.

PLMs such as ProtT5 [18], ESM-1b [19], ESM-1v [20] and ESM-[21] have been trained on a large corpus of raw protein sequences with the objective of predicting missing or masked amino acids given the context of the non-masked sequence. This results in a learned feature representation called an ‘embedding’ for each residue position in the protein sequence. The embeddings of these sequence-based pre-trained models have been shown to encode useful bio-physical information, such as residue conservation [22] amino acid hydrophobicity, protein structure class [18] and protein functional properties [23].

Even though these models were pre-trained without using evolutionary information, it has been shown that the methods achieve a similar performance to MSA-based models for various tasks while also reducing the computational cost.

Recent studies have used experimental data to evaluate the performance of PLMs in predicting the functional effects of variants [20,24]. However, to date, only one study (‘ESM variant’) has used a PLM to predict the clinical significance of a mutation [25]. ‘ESM variant’ uses the ESM-1b pre-trained PLM without requiring any supervised training. Given that ESM-1b was trained to predict the likelihood of each amino-acid type at each position, it is possible to use these likelihoods as a proxy for how well tolerated an amino-acid change is likely to be at the mutation site. ‘ESM variant’ constructs a pathogenicity score for a given mutation by using the ESM-1b likelihoods for the wildtype and mutant type amino acids at the mutated position **Fig 1**. The authors reported that their model outperformed EVE at variant pathogenicity prediction (AUC-ROC score = 0.905 *vs*. 0.885).

**Fig 1.**
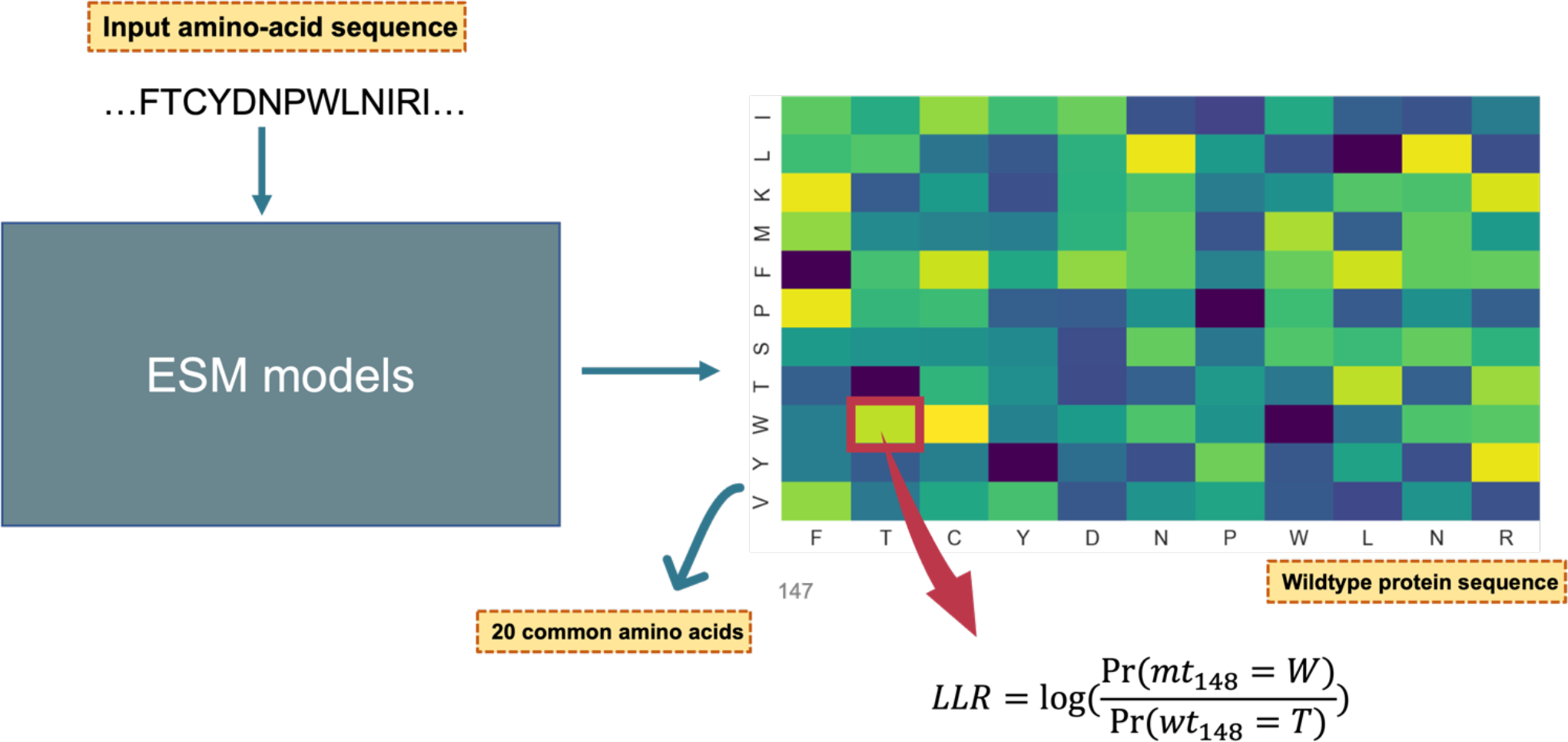
Illustration of how to calculate the log-likelihood ratio (LLR) from ESM models. The input for the PLM is an amino-acid sequence while the output is the log-likelihood ratio calculated based on the probabilities of the wildtype amino acid and the mutant amino acid occurring at the given position. **Note, the heatmap showing in this figure was artificially generated for the demonstration purpose. Pr**, Probability; wt_148_ = T, the148^th^ residue occurring at the input wildtype (**wt**) protein sequence is a Threonine (**T**); mt_148_ = W, the148^th^ residue was mutant type (**mt**) Tryptophan (**W**).

This study extends the research on PLMs by proposing a novel methodology and architecture. We conduct a comparative analysis of alternative PLM models including the recent ESM-1v and ESM-2 models which were trained on larger protein datasets. We compare performance of our model against EVE, ‘ESM variant’ and 3Cnet using the ClinVar dataset. Additionally, we investigate whether the predictors are prone to bias from Type 2 data circularity using two additional benchmarks: SwissvarFilteredMix and VaribenchSelectedPure.

Our model, VariPred, uses a novel twin-network PLM architecture combined with a trained classification module to achieve the highest classification performance on two variant classification benchmarks. The twin neural network framework (sometimes called a Siamese network) describes an approach where two comparable inputs are each passed through the same network. In our case, we pass the mutant and wildtype sequences through the PLM to generate embeddings for each residue position. Subsequently, we concatenate the two embeddings for the wildtype and mutant type residues that occur at the mutation position. These paired embeddings are used as input features to a lightweight feed-forward classification module which is trained on the labelled data. As a result of the transformer network architecture and the PLM’s pre-training objective, contextual information from the entire sequence is incorporated into the per-residue embedding. At the same time, selecting only the embeddings for the residues at the mutation position allows the classification module to focus on information which is specific to the mutation site.

## 2 Methods

### 2.1 Dataset preparation

Since ESM variant and EVE are both unsupervised learning models, they do not require a training dataset. To avoid having to retrain 3Cnet we opted to use the same train/test split as the 3Cnet authors. The only modifications that we made were to exclude the 3Cnet simulated data from our model’s training set and remove some variants from the test set which appear to have been inadvertently included by the 3Cnet authors in both the train and test sets.

#### 2.1.1 Training set

The training dataset used in 3Cnet consists of three parts: 1) clinical data stored in the clinical database ClinVar, 2) common missense variants retrieved from the population database GnomAD, and 3) a set of simulated pathogenic data generated by the 3Cnet authors. The simulated data are based on amino acid conservation determined from MSAs, built using sequences from the RefSeq database. We chose to exclude the simulated data from our training dataset and work with the subset of the 3Cnet training data sourced from ClinVar and GnomAD.

The ClinVar dataset used by 3Cnet was downloaded from the ClinVar database via the FTP link (version 2020-4). In total, 72,470 curated missense variants were selected according to the criteria of known molecular consequences and reliable review status. Specifically, only variants with the GRCh37 assembly version and labelled with ‘missense variants’ were collected, and those with unreliable review status, containing strings with ‘no assertion’, ‘Conflicting’, ‘no interpretation’, and ‘Uncertain’ were all excluded. Data labelled with ‘pathogenic’ or ‘likely pathogenic’ were all considered as pathogenic variants. Similarly, variants with any submission reported as either ‘benign’ or ‘likely benign’ were defined as neutral. After filtering out low-quality data, 72,470 variants (22,337 pathogenic and 50,133 benign) remained.

The GnomAD dataset prepared by the 3Cnet group (file downloaded using FTP: gnomad.exomes.r2.0.2.sites.vcf.gz) consists of 60,614 exome-derived variants. These variants have a minor allele frequency (MAF) higher than 0.1% and each was filtered by requiring a ‘PASS’ annotation, which ensures the quality of the variant, i.e. high confidence genotypes. Since these variants are found in the genome of supposedly healthy people, they are typically regarded as benign variants [26]. However, even though the 3Cnet authors regard variants with a MAF ≥ 0.1% as neutral, we cannot exclude the possibility that some of these variants have undetected (or partial penetrance) pathogenic effects.

Each datapoint included in these three datasets was annotated with a specific RefSeq NP code (protein record identifier in the protein sequence database) and the mutant information in the HGVSp term by the 3Cnet group, e.g. NP_689699.2:p.Gly56Ser. For each RefSeq NP code, the corresponding curated wildtype protein sequence was also provided by the 3Cnet group. For each variant in the dataset, the input for the model consists of both wild-type and mutant sequences, target mutated position, wildtype amino acid and the mutant amino acid. The final output of the model is a binary label, where 0 indicates that the mutation is benign and 1 indicates pathogenic.

#### 2.1.2 Test set

We noted that some variants (the same gene with the same mutation) were repeated between the 3Cnet train and test datasets. This problem arises from splitting the data based on variant information given in the HGVSp term. We found that some proteins annotated with different NP codes are in fact the same isoform, with the same wildtype protein sequence. To avoid having to retrain 3Cnet we chose to remove the 1,767 duplicated variants from the ClinVar test set and a further 900 variants which were duplicated between the GnomAD training dataset and the ClinVar test set.

As a result of removing these duplicates, the processed training dataset consists of 72,466 variants from ClinVar and 59,018 variants from GnomAD. In total 17% of variants were labelled as pathogenic. The test set is comprised of data from ClinVar only and consists of 21,125 entries with 45% of variants labelled as pathogenic (**Supplementary Table 1**). Here, we ensure that the training and test sets do not have any variants which are the same. Although all methods being benchmarked should ideally be tested using the same test set, for EVE the MSAs and predictions are not available for a significant proportion human genes, and thus we can only examine the performance of EVE on a subset. As a result, the test set for EVE shrinks from 21,125 to 4721, 52% of which were pathogenic, 28% Benign and 19% Uncertain (**Supplementary Table 2**).

#### 2.1.3 Testing for Type 2 data circularity bias

Grimm *et al*. [12] points out that effective benchmarking of clinical variant prediction can be confounded by circularity arising from overlap between the train and test sets. Type 1 data circularity arises when the same variant is included in the train and test set. Type 2 circularity arises from the same genes being included in the train and test set, even where the individual mutations are distinct. Type 2 data circularity bias is a particular problem where the data includes genes where labels are imbalanced towards one class. This scenario can give rise to predictors that ignore the specific details of the mutation merely recognising genes which are oversampled as pathogenic or benign in the training data. In order to assess the models’ propensity to overfit to genes in this way, we tested predictors using the SwissvarFilteredMix dataset and the VaribenchSelectedPure public benchmarks [27]. These datasets are used together to test whether performance is confounded by type 2 data circularity.

The SwissvarFilteredMix dataset consists of proteins with at least one type of label from each class. By contrast, in the VaribenchSelectedPure dataset each protein only has one type of variant class, either all benign or all pathogenic. If a model learns to predict based on characteristics of the gene and ignores the specifics of the variant, then it will typically show inflated performance on the VaribenchSelectedPure dataset while showing low performance on SwissvarFilteredMix. The 3Cnet and VariPred training set includes 67% of the genes that were in the VaribenchSelectedPure test set, although no variant was repeated between the train and test sets.

The VaribenchSelectedPure and SwissvarFilteredMix datasets contain information on chromosome number, base substitution position, reference nucleotide base, altered nucleotide base, Ensembl protein ID and the ground-truth label. Some sequences are not consistent with the mutation information, possibly because there has been a new isoform since the two benchmarks were generated in 2016.

Therefore, we annotated NM codes (mRNA record identifiers in the Nucleotide database) for each variant using the latest version of the ANNOVAR software [28], with the transcript-based annotation set for the RefSeq Gene (assembly version hg19; updated 2020-08-17 at UCSC). We then retrieved the corresponding protein isoform sequences (wildtype sequences) using the Entrez.efetch module included in Biopython (version 1.80) with Python 3.9. Using the original protein isoform sequences and the corresponding variant information, we generated the mutant sequences for each variant. This gave a SwissvarFilteredMix test set with 1153 benign variants and 1023 pathogenic variants, and a VaribenchSelectedPure set with 3629 benign variants and 2122 pathogenic variants.

Owing to the absence of some human genes in the EVE dataset, only 30% and 4.4% of variants in the SwissvarFilteredMix and VaribenchSelectedPure sets respectively, were annotated with the EVE binary pathogenicity label. Therefore, we could only keep 667 out of 2176 variants from SwissvarFilteredMix (278 benign *vs*. 238 pathogenic) and 254 variants from the VaribenchSelectedPure set (127 benign *vs*. 72 pathogenic). Moreover, 151 out of 667 variants in the SwissFilteredMix and 55 out of 254 variants in the VaribenchSelectedPure set were labelled as ‘uncertain’. As a result, this dataset is much smaller than for the other methods; nonetheless, we still evaluated the performance of EVE for comparison.

As with EVE, there are also problems with 3Cnet in generating features for some variants from these two benchmark datasets. To predict a variant’s pathogenicity with 3Cnet requires three components: the HGVSp term including the NP code and the mutation information, the NP code corresponding to the wildtype sequence, and 85 biological features retrieved from the SNVBox database [29]. However, this information is not recorded in the SNVBox database for some of the variants. Consequently, 3Cnet is not able to give a prediction for these variants. We therefore dropped these variants, leaving 1742 variants in the SwissvarFilteredMix and 5159 variants in VaribenchSelectedPure test sets for the evaluation of 3Cnet.

### 2.2 Feature extraction and model setup

In order to identify the most suitable PLM for differentiating pathogenic variants from benign, we tested the most widely used pre-trained models including ESM-1b, ESM-1v and ESM-2. ESM-2 has several versions with different parameter sizes, ranging from 8×10^6^ to 15×10^9^. According to a previous study, the performance of the model does not increase with model size, and models with 650×10^6^ parameters appear to have the best ability to extract per-residue features [30]. Therefore, we chose ESM-2 with 650×10^6^ parameters for our analyses.

#### 2.2.1 Extract embeddings by PLMs

ESM-1b and ESM-1v are BERT-style encoder-based Transformers, which limit the input length to 1022 amino acids. ESM-2 does not have this sequence length limitation, but using longer sequences is computationally prohibitive. Moreover, the Rotary Position Embedding strategy used in ESM-2 only considers the word embeddings and their neighbours, limiting any advantage of larger windows. Therefore, we designed a sequence truncation strategy which is consistent with such encoding methods and limits the maximum length to 1022 in all 3 models.

The official ESM tokenizer package pads the length of shorter sequences to 1022 internally, but transforms the length back to the true sequence length during data processing. For sequences longer than 1022, if the mutation is within 1022 residues of either the N-terminus or the C-terminus, 1022 residues counting from the end were retained; if the mutation index occurs more than 1022 residues from both termini, 510 neighbours from the N-terminal side and 511 from the C-terminal side of the mutated residue were selected, resulting in sequences having 1022 residues.

Transformer-based PLMs provide features in two forms: the probability of each amino acid type occurring at each position in the sequence, and a dense vector-embedded representation of each position in the sequence. Owing to the self-attention mechanism of the Transformer architecture, the embedding can incorporate contextual information from the entire 1022 residue window. For each mutated position we extracted the log likelihood ratio (LLR) and the embedded representation of the wild-type and mutant type residue.

The LLR was calculated using the ESM likelihood of the mutant and wildtype amino acid at the target position conditioned on the model receiving the wildtype sequence as input, using the formula shown in **Fig. 1**. PLMs are pre-trained using a masked language modelling objective. During the pre-training, 15% of residues were randomly masked out from the input sequences, and the model predicts which amino acid type is most likely to be present at each masked position.

Amino acids which frequently occur at a target position, typically have a comparatively high likelihood. Thus, if the mutant type’s likelihood is significantly lower than the wild type, this serves as an indicator that the mutation is problematic, while mutant residues with high likelihoods typically have similar physiochemical properties and are therefore less likely to affect protein stability or function. The ‘ESM variant’ method [25], which uses the LLR generated by ESM-1b to discriminate variant pathogenicity, claimed that an LLR threshold of -7.5 is sufficient to detect pathogenic variants. Therefore, in this study, we also applied -7.5 as the LLR threshold for ESM-1v and ESM-2. The ESM models evaluated in this study use word embedding dimensions set to 1280 to represent each position in the sequence.

The schema of data processing and model generation is given in **Fig. 2**. To obtain the embeddings of each sequence pair (wildtype and mutant protein sequences), all sequence pairs were fed into the PLM. For example, if a wildtype sequence consists of 100 amino acids, two embedding matrices (one for amino acids in the wildtype sequence, the other for amino acids in the mutant sequence) with dimensions 100 × 1280 would be generated (**Fig. 2A**).

**Fig 2.**
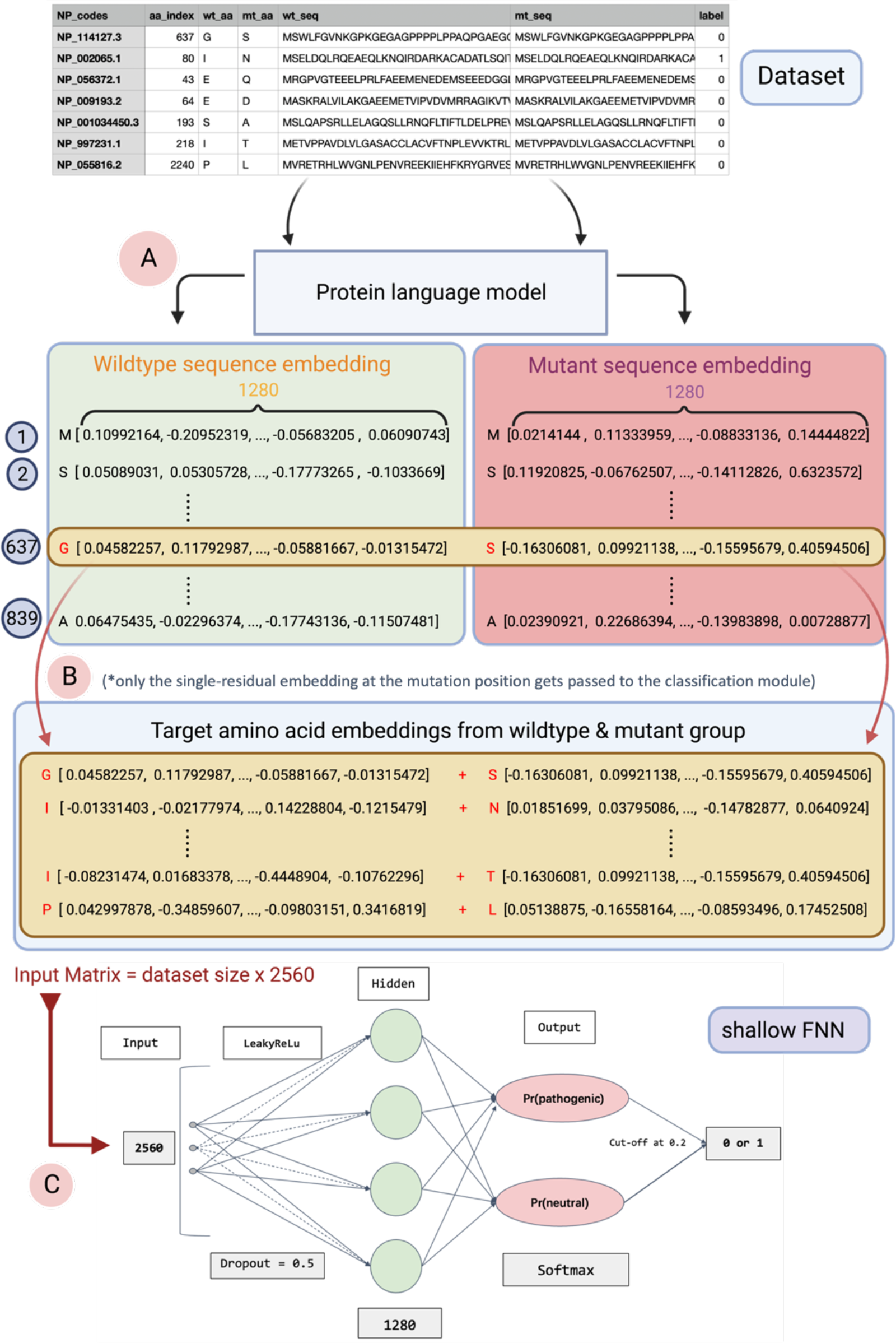
Schema of workflow for training VariPred. A) In the first step, each wild-type protein sequence and the corresponding mutant protein sequence are fed into the PLM separately. The PLM generates a per-residue embedding for each amino acid. The output is the matrix of sequence embedding, with dimensions sequence length x embedding dimension. B) Only the embeddings of the amino acids at the mutated position are used and joined giving an embedding dimension of 2560. The concatenated embeddings for each observation are combined to give an embedding matrix with dimensions dataset size x 2560. C. The embedding matrix is fed as the input into a Feedforward Neural Network (FNN), and two probabilities are then output identifying if the given variant belongs to the pathogenic or benign group. **Note that if the LLR feature is appended the input matrix is dataset size X 2561**.

We hypothesised that the embedding of the target amino acid at the mutation position would be the most informative. Therefore, we only took the embedding of the amino acid at the mutated position from both wildtype and mutant sequences. These two embeddings were then concatenated horizontally such that each data entry is represented as a vector with dimensions 1 × 2560. Feeding the training dataset (192575 entries) into the ESM-1b pre-trained model will generate a wildtype-mutant concatenated amino acid embedding representation matrix with a size of (192575 × 2560) (**Fig. 2B**). These embeddings are expected to capture fundamental biological features, related to protein function or structural stability.

To investigate whether combining LLR and embeddings would increase performance, we appended the LLR to the last column of the embedding matrix, which increased the dimension from 2560 to 2561 (**Fig. 2C**).

#### 2.2.2 Feed-Forward Neural Network

We created a classification module by including a shallow feed forward neural network (FNN) as the decoder/classifier for the PLM. This was trained on the class labels without updating parameters in the PLM. During the hyperparameter tuning process, we tried increasing the depth of the FNN as well as trying multiple sets of learning rates and drop-out rates. The final FNN, which gave optimal performance, consists of one hidden layer, a LeakyReLu activation function, and one output layer with the dropout rate set at 0.5 and learning rate set at 0.0001 (**Fig. 3C**). The input layer of the feed forward neural network has 2560 nodes (2561 if LLR is appended), while the hidden layer contains 1280 nodes, and the output consists of 2 nodes with a SoftMax layer to ensure the output probabilities sum to 1 for binary classification of benign/pathogenic. Only the pathogenic output node was considered and a value of 0.2 was selected as a threshold for predicting the pathogenic class. This low threshold was selected because the dataset is highly skewed towards neutral variants and this value optimizes the MCC on the training data.

**Fig 3.**
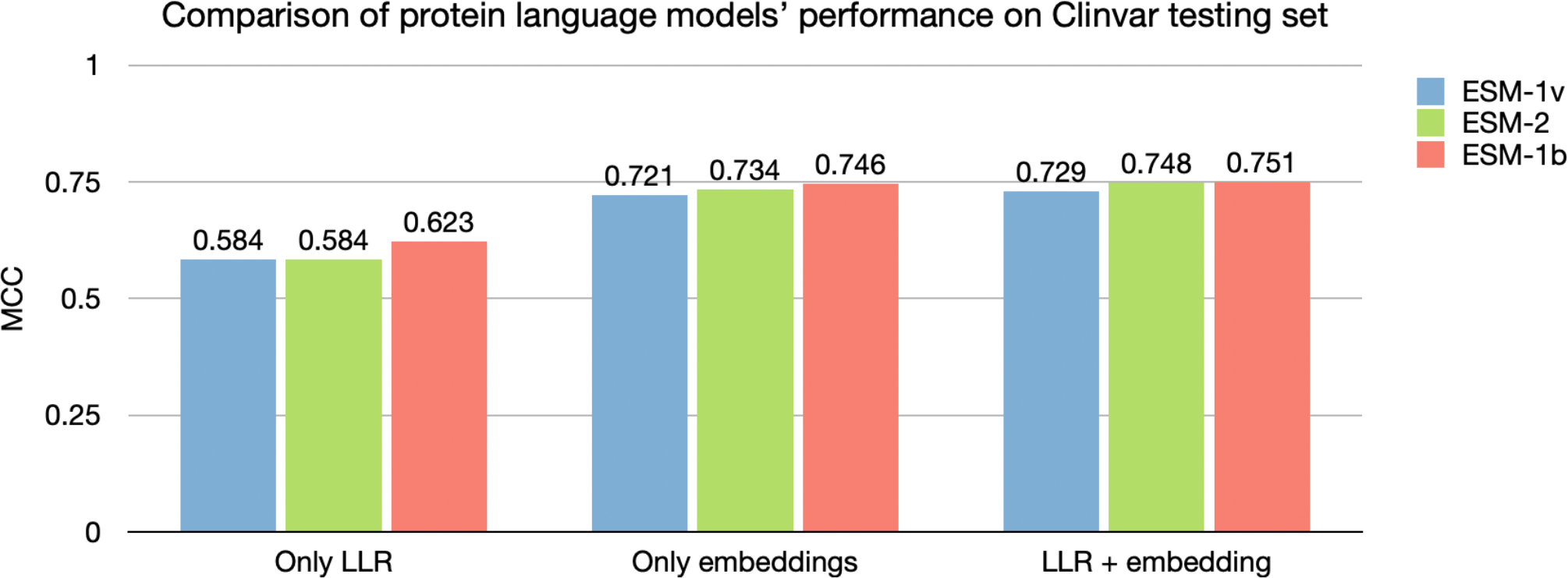
Comparison between models under different testing situations. Comparison of the LLR and embedding features for three protein language models. The baseline is using ‘Only LLR’ to predict pathogenicity of variants; For ‘Only embeddings’ we used amino acid embeddings as input to a shallow FNN to predict the pathogenicity of variants; ‘LLR + embeddings’ concatenates the LLR feature as the last column of the amino acid embedding matrix, and then performs variant classification by using this extended matrix as input to the FNN.

### 2.3 Evaluation metrics

Accuracy, Precision-Recall, F1-score, MCC (Matthews correlation coefficient) and AUC-ROC (area under curve of the receiver operating characteristic) are some of the most popular metrics for evaluating binary classifiers. Since MCC takes all outcomes (true and false positives and negatives) into account (Equation 1), it is less sensitive to class imbalance and is also more informative about the classifier’s performance at a given threshold [31]. In contrast, other measures are more sensitive to imbalance [32] and the AUC-ROC gives a view of the overall performance of a classifier (across a range of thresholds) rather than the actual performance in a classification problem.

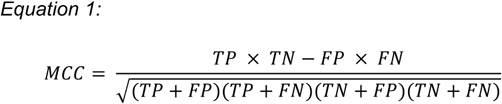

Therefore, we applied MCC as the main metric to measure the performance of predictors studied in this research, while using AUC-ROC as an auxiliary indicator.

## 3 Results

### 3.1 Comparing the performance of LLR, embeddings and LLR + embeddings

The ‘ESM variant’ method [25] only uses the LLR feature generated from ESM-1b to predict the clinical significance of missense variants. Here we evaluate the performance for this task, using LLRs generated by two other PLMs: ESM-1v and ESM-2.

We first tested models using the ClinVar test set. For PLMs which only use the LLR threshold for prediction, ‘ESM variant’ (using ESM-1b) has the best predictive performance, with an MCC score of 0.623, followed by ESM-2 (0.5837) and ESM-1v (0.584) (**Fig. 3**).

In addition to using LLR threshold to differentiate pathogenic variants from benign, our VariPred predictor uses a shallow FNN trained on the PLM embeddings for the wildtype and mutant sequence. We observe that this approach significantly improves the performance of all protein language models. ESM1b, remains the best performing model, with an MCC score of 0.746, followed by ESM-2 (0.734) and ESM1v (0.721) (**Fig. 3**).

When we combine the LLR together with the embeddings, the performance of all models improved further. ESM-1b still has the best performance with an MCC score of 0.751, ESM-2 scored 0.748, and ESM-1v achieved 0.728 (**Fig. 3)**. Therefore, in the following experiments to compare the performance with other tools, we chose ESM-1b as the feature extractor for our model, VariPred. We trained our model with both embeddings and LLRs as input.

### 3.2 Comparison between models using the ClinVar test set

To evaluate VariPred’s performance on clinical data, we compared the performance of VariPred with other tools on the ClinVar test set. Since EVE also classifies variants with a third label “Uncertain”, we calculated two MCC scores for EVE. In the first case, we considered variants labelled as “Uncertain” as pathogenic and then as benign, and calculated the mean of the MCCs resulting from these two scenarios as the first score for EVE (EVE-avg). In the second case, we simply ignored “Uncertain” predictions, and calculate the MCC score of EVE with the remaining data (EVE-ign). In this case, 20% of the data were dropped and only 3469 variants remained for the evaluation.

Comparing against other methods, VariPred has the best performance with an MCC of 0.751. 3Cnet is closest to matching the performance of VariPred with an MCC of 0.690, followed by EVE-ign (MCC=0.673) and ‘ESM variant’ (MCC=0.620), while EVE-avg has the lowest performance with an MCC of 0.545 (**Fig. 4**). However, VariPred only requires protein sequence information as input, while 3Cnet requires features including MSAs, amino acid physicochemical properties, and protein features such as motifs or active sites.

**Fig 4.**
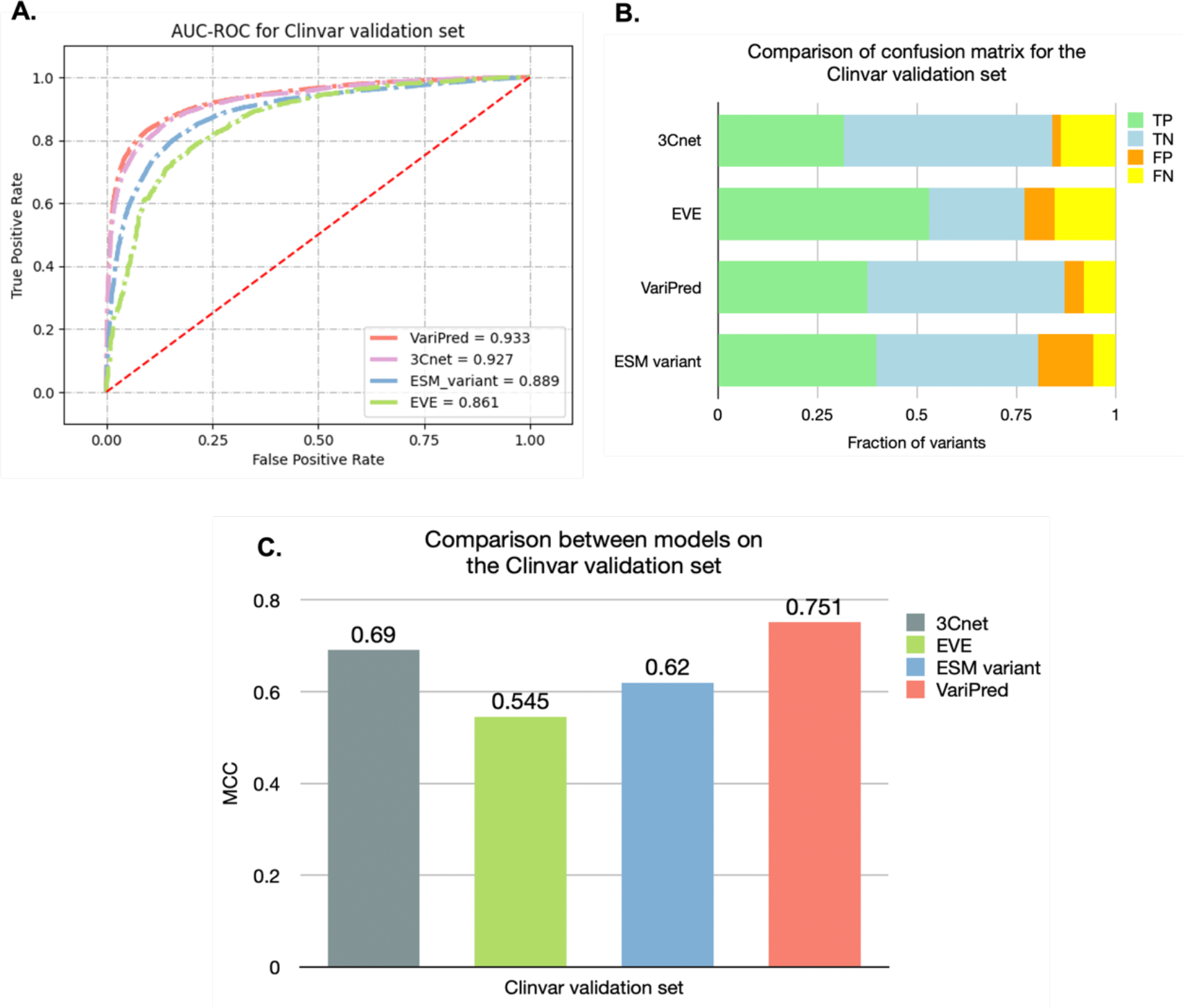
Comparing the performance of pathogenicity predictors using the ClinVar validation set. A) AUC-ROC curve plot for the four predictors. B) The Confusion matrix comparison for predictors. Note that EVE has a smaller test set size due to the problems with data availability. Consequently, the distribution of actual positives and actual negatives is different from the other predictors. All overlaps of mutant protein sequences, between the training set and this test set have been removed. C) MCC score for the predictors being tested in this study.

### 3.3 Type 2 data circularity problem test

We compared all predictors including the three PLMs with only LLR (as the baseline) and the combination of LLR and embedding features with two public benchmarks, the SwissvarFilteredMix and VaribenchSelectedPure test sets, to evaluate if any of the models are affected by the Type 2 data circularity problem.

In this evaluation, 3Cnet has the highest accuracy in VaribenchSelectedPure, and is the only model with higher performance in VaribenchSelectedPure (MCC=0.403) than SwissvarFilteredMix set (MCC=0.161) (**Fig. 5**). This suggests that 3Cnet is affected by the Type 2 data circularity problem where the model is learning features of the gene and ignoring the specifics of the variant. In contrast, our model, VariPred, has the second highest performance in SwissvarFilteredMix (MCC=0.440) but a lower performance in VaribenchSelectedPure (MCC=0.240) (**Fig. 5**), indicating that VariPred is not confounded by the Type 2 data circularity problem.

**Fig 5.**
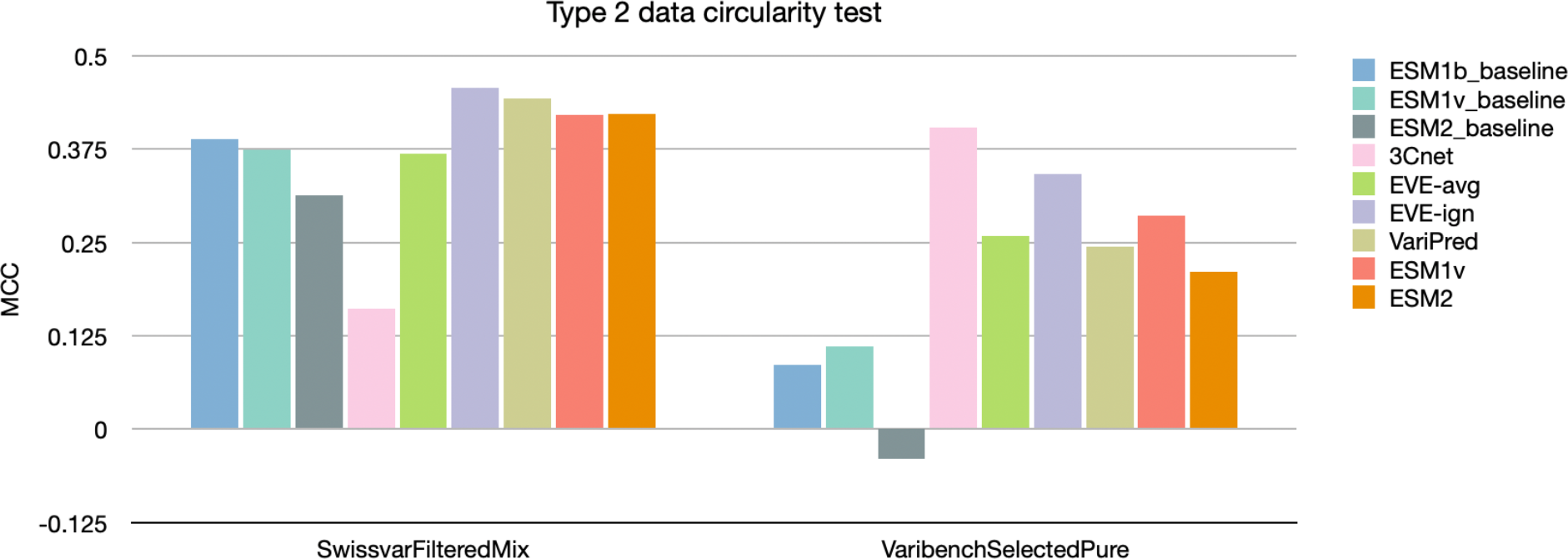
Type 2 data circularity problem test. Predictors labelled with baseline only use LLR to classify variants. The ESM-1v and ESM-2 methods only use embedding features as input and have further supervised training using the training set mentioned in the Methods.

We evaluated EVE (both -ign and -avg) on the SwissvarFilteredMix and VaribenchSelectedPure test sets with the publicly available pre-computed data. The MCC for EVE is 0.349 when regarding all variants labelled as Uncertain as pathogenic, and is 0.387 when considering them as neutral, giving a mean score (EVE-avg) of 0.368. On the other hand, EVE-ign has the best performance in this test set, with an MCC of 0.450, but no prediction is made for approximately 20% of the data. We removed variants with no pre-computed EVE score from VaribenchSelectedPure, giving a small dataset of 254 (127 benign *vs*. 72 pathogenic *vs*. 55 uncertain). EVE-avg with VaribenchSelectedPure gave an MCC of 0.259, while EVE-ign gave a higher MCC of 0.325, but with only 199 variants included in the analysis. Both EVE-avg and EVE-ign give MCCs lower than that on the SwissvarFilteredMix test set (**Fig. 5**) suggesting there is no issue with Type 2 data circularity, although this has been assessed using a much smaller dataset than for the other predictors.

## 4 Discussion

We tested three different PLMs (ESM-1b, ESM-1v, ESM-2) and showed that ESM-1b was the best predictor for pathogenicity of single-position missense variants. Using the ClinVar test set, our VariPred predictor which combined LLR and residue embeddings generated by ESM-1b has the best performance achieving an MCC of 0.751 and AUC-ROC of 0.933 without using any additional biological features and not being confounded by Type 2 data circularity.

In principle, since ESM-1v and ESM-2 were pre-trained using a much larger protein sequence dataset, they should have a broader view on the mutability landscape of proteins than ESM-1b. The ESM-1v and ESM-2 authors state that both ESM-1v and ESM-2 are sufficient to conduct the missense mutation pathogenicity prediction without any further training [20,30]. Nonetheless, we observed better performance on the ESM-1b model. We speculate that this may be a result of the ESM-1b pre-training dataset being more closely aligned with the relatively narrow set of (human only) proteins that are included in ClinVar.

Reports suggest that for predicting the functional (rather than clinical) effects of variants, which are in the form of a continuous scalar value, ESM1-v and ESM-2 have a better performance [20,30]. However, recent comments suggest that ESM-1b performs better in some other tasks, such as structure prediction [33]. In this study, we showed that ESM-1b outperforms two other state-of-the-art predictors in predicting the binary clinical significance of missense variants.

Although 3Cnet has a similar performance to VariPred, it has a better performance on the VaribenchSelectPure dataset compared with the SwissvarfilteredMix set (see **Fig. 5**). This indicates that 3Cnet is affected by the Type 2 data circularity problem and may be more prone to biased predictions arising from the composition of the training data.

Besides the leading performance of VariPred in predicting the clinical significance of missense variations, VariPred is more efficient in facilitating a high-throughput prediction of the pathogenicity of variants in humans as it has minimal dependencies and only requires the sequence as input.

Many human genes are not included in the MSAs provided by EVE or in the pre-computed predictions, therefore we have not evaluated its performance on genes which have shallow MSAs. We note that collecting the MSAs for each protein adds a large computation and memory cost and computing the EVE score for each variant is also highly computationally intensive.

Preparing the input features for 3Cnet is a difficult task. 3Cnet relies on features based on 85 biophysical properties retrieved from the SNVBox database. However, not all variants can be mapped with features from the SNVBox database as it has not been updated since 2011, resulting in missing sequences and problems with changed RefSeq IDs. This may lead to uncertainty in the consistency between the retrieved feature and the data entry. Additionally, only the NP codes which are included in the provided transcript ID list can be transformed into a 3Cnet prediction dataset.

In comparison to EVE and 3Cnet, VariPred requires only the most fundamental information for each data entry: the wildtype protein sequence and mutation information including ***which*** residue is being mutated at ***which position*** in the wildtype sequence into ***which*** mutant amino acid. Without the need for further dataset preparation, such as MSA construction or feature retrieval, making predictions on a dataset of 5000 variants take 30 minutes with a 12GB GPU, such as the Nvidia GTX 1080Ti.

We note that performance on the SwissvarFilteredMix testing set was lower than performance on ClinVar for all models. Comparing this dataset with the latest ClinVar dataset (2022-12) we found that 17% of entries have an inconsistent label (e.g. ‘uncertain’ in SwissVarFilteredMix versus ‘positive’ in ClinVar) and 81% do not have a label in the ClinVar dataset, whilst 2% of variants have the opposite label. This could explain why the performance of all predictors drops in the SwissvarFilteredMix set, as the inconsistent data would reduce the evaluated prediction performance of the predictors. This could be a result of different criteria for labelling (e.g. how partial penetrance variants are classified) or labels changing due to newly arisen evidence.

Inconclusive and contradictory pathogenicity labels are an argument in favour of unsupervised methods such as EVE and ‘ESM variant’. Even though we observed lower performance when compared with supervised methods such as VariPred and 3Cnet, it is important to note that the unsupervised methods are not prone to bias introduced by training dataset selection and labelling issues.

Several authors have suggested that predictors may have better performance on variants of a specific gene or disease [27,34]. In the future, we will evaluate VariPred’s performance on specific genes associated with various diseases as well as differential pathogenicity prediction – i.e. predicting different pathogenic phenotypes caused by mutations in the same protein [35].

Currently, VariPred only uses sequence information to predict pathogenicity. In the future we will evaluate the effect of including structural information in the predictor. A similar strategy has been implemented for aiding engineering of enzymes by directed evolution and for aiding protein design [36,37], but has not yet been explored in the prediction of the clinical significance of missense mutations. Incorporating both sequence and structural information is likely to improve VariPred’s ability to classify missense variants.

Applying a more biologically meaningful data augmentation strategy may add more diversity into the training set. Conservation information is one of the most powerful features for predicting protein stability and functional effects [38]. In the study of 3Cnet, the artificial pathogenic-like variants were generated simply by considering the amino acid frequency and the number of gaps. However, a good conservation scoring scheme depends on multiple components, of which the most important include amino acid frequency, residue similarity (biophysical properties), sequence similarity (considering sequence redundancy and MSA depth), the number of gaps in the MSA, and the concept of ‘compensated pathogenic mutations’ (CPDs), which refers to mutations occurring in different species that are tolerated because of compensating mutations [39]. Therefore, in the future, it may be worth investigating whether such a combination of data augmentation and synthetic data strategies can further improve the performance of VariPred.

In summary, VariPred only requires the native and mutated sequence and, using protein language model encoding, is able to outperform state-of-the-art methods that use features including structural information and multiple sequence alignments.

#### Keypoints

Our main contributions are summarised as follow:

- VariPred is a new model for predicting the clinical significance of missense mutations. We combine the ESM-1b protein language model with a twin-network architecture that has not been applied within this field previously.
- We demonstrate that this novel approach outperforms existing state of the art methods on multiple benchmark datasets.
- The model only requires the protein sequence as input, has minimal dependencies and unlike competing methods it does not require MSAs. This makes it faster and easier to use.

## Code availability

All code required to reproduce the model and analysis in this study are available at https://github.com/wlin16/VariPred.git.

## Acknowledgements

The authors thank Jinyuan Sun, Nicola Bordin, Ian Sillitoe, Marcia Hasenahuer, Shi-Yuan Tong for help with Bioinformatics tools including HHblits, MMseqs, Blastp, Jackhmmer and Scorecons. The authors thank David Gregory for help with implementing the SNVBox database and Jin Dai for help with implementing protein language models.

