## Supplementary Table for "VariPred: Enhancing Pathogenicity Prediction of Missense Variants Using Protein Language Models"

**Supplementary Table 1. The number of pathogenic and benign variants of each set included in the training set before and after dropping the duplicates within itself.**

| Training set |  |  | Clinvar | Gnomad |
| --- | --- | --- | --- | --- |
|  | Before duplicates drop | Patho | 22337 | 0 |
|  |  | Benign | 50133 | 60614 |
|  | After duplicates drop | Patho | 22334 | 0 |
|  |  | Benign | 50132 | 59918 |
|  | After overlap drop | Patho | 22334 | 0 |
|  |  | Benign | 50132 | 59018 |
|  | After Swissvar | Patho | 22254 | |
|  |  | Benign | 108899 | |
|  | After Varibench | Patho | 22235 | |
|  |  | Benign | 108449 | |

**Supplementary note for Table 1:** We first dropped the duplicates within the two sets used as the training set. Then, we dropped the overlaps between these two sets. Subsequently, we dropped the overlaps between the SwissFilteredMix set (Swissvar) and the training set from the training set first, and finally removed the same variants between the VaribenchSelectedPure set (Varibench) and the training set from the training set. The part highlighted in yellow indicates the amount of data in the dataset was not changed in that stage.

**Supplementary Table 2. The number of pathogenic and benign variants of each test set before and after dropping the duplicates within itself, dropping the overlaps between the training set, and dropping the data which are not available for a prediction by the tool.**

| Test set |  |  | Clinvar | SwissVar | VaribenchPure |
| --- | --- | --- | --- | --- | --- |
|  | Before duplicates drop | Patho | 12041 | 1153 | 3629 |
|  |  | Benign | 10262 | 1023 | 2122 |
|  | After duplicates drop | Patho | 11930 | 1153 | 3629 |
|  |  | Benign | 9993 | 1023 | 2122 |
|  | After overlaps with training set | Patho | 11514 | 1153 | 3629 |
|  |  | Benign | 9611 | 1023 | 2122 |
|  | After 3Cnet | Patho | 11514 | 776 | 2049 |
|  |  | Benign | 9611 | 966 | 3110 |
|  | After EVE | Patho | 2459 | 278 | 72 |
|  |  | Benign | 1343 | 238 | 127 |
|  |  | Uncertain | 919 | 151 | 55 |

**Supplymentary note for Table 2:** We first dropped the duplicates within each test set. In this stage, since the SwissFilteredMix set (Swissvar) and the VaribenchSelectedPure set (Varibench) do not have any duplicates, the size of both sets remains still. In the next stage, again, since we removed the duplicates between the training and the SwissFilteredMix set and the VaribenchSelectedPure set from the training set, the size of these two sets was not changed. In the following stage, due to the limitation of 3Cnet and EVE, of which the impacts of some variants are not available to be predicted, we removed these variants from the test sets. For the ClinVar test set, since it was originally provided by the 3Cnet, thus, the effects of all variants (21156) are available to be predicted by 3Cnet (highlighted in yellow indicates an unchanged dataset size). However, some genes in the ClinVar test set are not available for a prediction by EVE. Thus, when testing the effects of variants in the ClinVar test set by EVE, we can only obtain the prediction results of 4721 variants.
